# Protein Model Refinement for Cryo-EM Maps Using DAQ score

**DOI:** 10.1101/2022.08.23.505034

**Authors:** Genki Terashi, Xiao Wang, Daisuke Kihara

## Abstract

As more protein structure models have been determined from cryo-electron microscopy (cryo-EM) density maps, establishing how to evaluate the model accuracy and how to correct models in case they contain errors is becoming crucial to ensuring the quality of structure models deposited to the public database, PDB. Here, we present a new protocol for evaluating a protein model built from a cryo-EM map and for applying local structure refinement in case the model has potential errors. Model evaluation is performed with a deep learning-based model-local map assessment score, DAQ, which we developed recently. Then, the subsequent local refinement is performed by a modified procedure of AlphaFold2, where we provide a trimmed template and trimmed multiple sequence alignment as input to control which structure regions to refine while leaving other more confident regions in the model intact. A benchmark study showed that our protocol, DAQ-refine, consistently improves low-quality regions of initial models. Among about 20 refined models generated for an initial structure, DAQ score was able to identify most accurate models. The observed improvements by DAQ-refine were on average larger than other existing methods.

## 1. Introduction

Cryo-electron microscopy (cryo-EM) has become a core technique for determining the three-dimensional structure of biomolecules. Technical advancement of cryo-EM occurred a few years ago, which made it possible to achieve near-atomic resolution in determining density maps (Yip *et al*., 2020). Cryo-EM has unique strengths including the ability to determine macromolecular structures, such as large complexes or membrane proteins, that are often difficult to crystallize for X-ray crystallography (Cheng, 2018). As an increasing number of protein structures are being determined by cryo-EM, however, it has been observed that a substantial number of structure models determined with cryo-EM and deposited in PDB (Berman *et al*., 2000) have potential errors (Terashi *et al*., 2022). Errors include cases where modeled local conformation itself is wrong and situations where misassignment of amino acids occurred with otherwise correct main-chain conformation.

Structure modeling is intrinsically difficult for map regions where the local resolution is low. Also, errors often occur when assigning sequences along a long helix as amino acids in the sequence region all have high helix propensity. As the “resolution revolution” of cryo-EM (Yip *et al*., 2020) has opened the door of structural analysis to many inexperienced users who are attempting to build atomic models into maps of moderate resolution, many mistakes may have been introduced in structure models. Therefore, a reliable protocol is needed that can correctly identify and fix regions in models that are built from a cryo-EM map.

The cryo-EM modeling community recognizes the importance of model validation as it was a focus of discussions in the EM Modelling Challenge held in 2019 (Lawson *et al*., 2021). Various model validation methods have been proposed in the past. Validation methods can be categorized mainly into two types, map-model scores and model-coordinate scores. The former, map-model scores, measure the consistency of the map and a protein model, which include the atom inclusion score (Joseph *et al*., 2017), EMRinger (Barad *et al*., 2015), Q-score (Pintilie *et al*., 2020), and map correlation score (Joseph *et al*., 2017, Joseph *et al*., 2022). The other category, model-coordinates scores, such as MolProbity (Chen *et al*., 2010) and CaBLAM (Prisant *et al*., 2020), identify stereochemical outliers in a protein model. Recently, we developed a novel validation method, Deep learning based Amino acid-wise model Quality (DAQ) score, which uses deep learning to capture local density features to assess the likelihood that each residue in a model is correct. It is demonstrated that DAQ score was very effective in finding the positions in the structure model where amino acid assignments to a local density are likely to be incorrect (Terashi *et al*., 2022).

Following our development of DAQ score, here we address the next step – how to refine identified incorrect local regions in a structure model. The local refinement protocol, DAQ-refine, starts by identifying potential incorrect regions in a protein model using DAQ-score. Then, it remodels the local regions using AlphaFold2 (AF2), the protein structure prediction method (Jumper, Evans, Pritzel, Green, Figurnov, Ronneberger, Tunyasuvunakool, Bates, Zidek, Potapenko, Bridgland, Meyer, Kohl, Ballard, Cowie, Romera-Paredes, Nikolov, Jain, Adler, Back, Petersen, Reiman, Clancy, Zielinski, Steinegger, Pacholska, Berghammer, Bodenstein, *et al*., 2021, Jumper, Evans, Pritzel, Green, Figurnov, Ronneberger, Tunyasuvunakool, Bates, Zidek, Potapenko, Bridgland, Meyer, Kohl, Ballard, Cowie, Romera-Paredes, Nikolov, Jain, Adler, Back, Petersen, Reiman, Clancy, Zielinski, Steinegger, Pacholska, Berghammer, Silver, *et al*., 2021) that achieved significantly high accuracy in the 14^th^ Critical Assessment of Structure Prediction (CASP14), a community-wide protein structure prediction experiment (Kryshtafovych *et al*., 2021). Although AF2 produces a highly accurate model from the protein sequence information in many cases, there are several reported limitations (Aderinwale *et al*., 2022, Jones & Thornton, 2022). Among the known limitations, the most relevant issue for this work is that a predicted structure from AF2 is built solely based on the sequence and is occasionally different from the conformation in a particular experimental structure and condition, such as in a complex determined by cryo-EM (Zhou *et al*., 2020, Heo & Feig, 2022, Del Alamo *et al*., 2022). Thus, in the current refinement protocol, instead of running AF2 as it is we attempt to keep the confident regions in the initial protein model intact and to only remodel low-confident regions by AF2. This is achieved by following a recently proposed protocol with AF2, where a partial structure of the target protein is provided as a template to AF2 (Heo & Feig, 2022).

The current protocol is different and complementary to the protein structure modeling protocol that Terwilliger et al. proposed in the framework of the Phenix modeling package (Terwilliger *et al*., 2021). Their protocol, *dock_and_rebuild*, employs AF2 and the Phenix model fitting iteratively to model the entire protein structure model for a density map. In contrast to their approach, the protocol we discuss here is about correcting local regions in a structure model that was already built by some modeling tool.

We applied DAQ-refine to 13 PDB deposited models and corresponding cryo-EM maps reconstructed at 3.20-4.35 Å resolution. For 11 targets (85%), our local refinement protocol achieved the highest GDT-HA model assessment score (Kopp *et al*., 2007) among four other model refinement methods. Detailed analysis shows that the final model selection by DAQ (AA) score and the use of customized templates contributed to the high performance of the proposed method. The result demonstrated that DAQ-refine is able to identify misaligned regions in the initial models and refine those regions in an automated fashion.

## 2. Material and Methods

We illustrate the overall protocol of DAQ-refine in Fig. 1. It consists of five steps: (1) Initial model evaluation by DAQ; (2) Generating multiple sequence alignments (MSAs) and a template model as input for AF2; (3) model building with AF2 with the customized input data. (4) model refinement with Rosetta Relax; and (5) model selection with DAQ score. We describe each step more in details in subsequent subsections.

**Figure 1.**
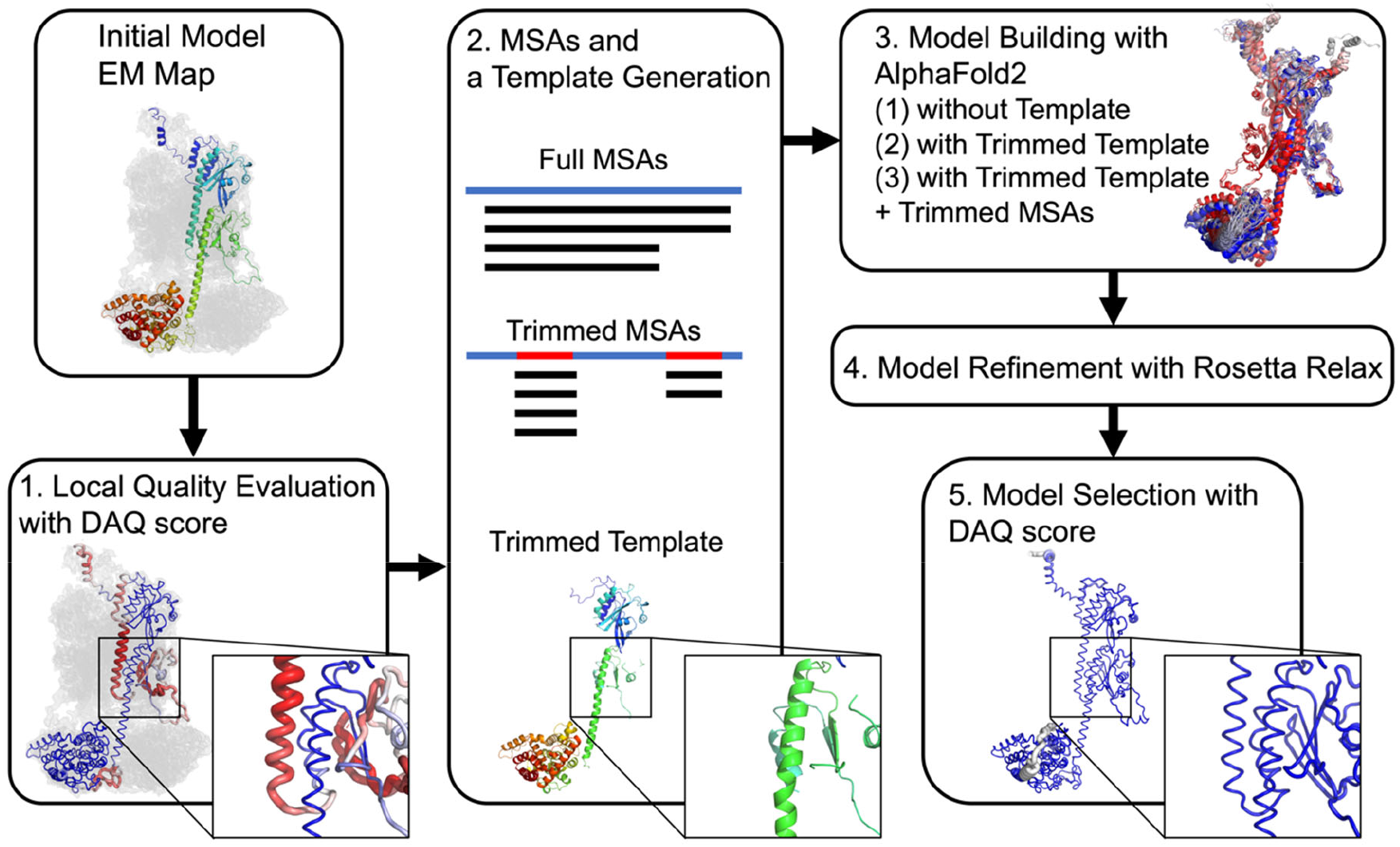
Overview of the DAQ-refine protocol. (1) The initial model evaluation with DAQ. The initially deposited model for PDB: 7JSN chain A, which was derived from the EM map (EMD-22458, 3.2 Å resolution) was used as an example. DAQ(AA) scores along the model are shown in a color scale from red (DAQ(AA) < −1.0) to blue (DAQ(AA) > 1.0). The magnified box highlights regions (residue His230 - Glu256) where DAQ(AA) is negative and thus highly likely to be incorrect. (2) MSA and template model generation. Full MSAs are computed by MMseq2 in ColabFold. Trimmed MSAs are generated by masking alignment data corresponding to the position in the full MSAs where the DAQ(AA) score is positive. The trimmed template is generated by removing residue positions where DAQ(AA) score is negative or zero from the initial model. (3) Model building by AlpfaFold2. Three strategies (AF2 with full MSAs, AF2 with full MSAs + trimmed template, and AF2 with trimmed MSAs + trimmed template) are performed. (4) Models are refined with Rosetta Relax in the EM map. (5) Finally, the top-ranked model by DAQ(AA) score is selected as the final model.

### 2.1. Initial model quality evaluation by DAQ

The first step in the protocol is to evaluate the model with DAQ score. DAQ score quantifies how well residues in an atomic model agree with density features detected by deep learning. Local density features of an EM map are captured by deep learning: The input EM map is scanned with a box 11*11*11 Å^3^ in size with an interval of 1 Å along the three coordinates axes directions and a trained deep neural network outputs probabilities that the box contains 20 amino acid types at the center position the box (Terashi *et al*., 2022).

The DAQ(AA) of a residue position *i* in a protein model is defined as

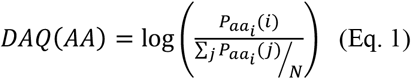

where *P*_*aai*_*(i)* is the computed probability of amino-acid type *aa*_*i*_ currently assigned at the position *i* by deep learning, which is normalized by the average value among all atom positions *j* in the model. *N* is the total number of atoms in the model. As the equation shows, if the predicted probability value for the position *i* is higher than the average, then DAQ(AA) becomes positive. If DAQ(AA) is below - 0.5 for an amino acid position, it is highly likely that the amino acid does not fit to the local density and worth attention. After DAQ(AA) was computed for all amino acids in a model, they were averaged by a sliding window of 19 residues along the protein sequence to better detect a local sequence shift in a model (Terashi *et al*., 2022).

DAQ score has two other types, probabilities of having a Cα atom, DAQ(Cα), and three types of secondary structures, DAQ(SS), in a scanning box. Here, we use only DAQ(AA) since it detects sequence assignment shifts and wrong conformation in a structure model, which are usually two common errors in protein structure modeling.

### 2.2. Running AlphaFold2 with modified MSAs and template

Detected low-confidence regions by DAQ-score in the initial model were refined with AF2. In this local refinement, we want to only refine the local regions but keep confident regions in the model untouched. A regular AF2 run is not suitable for this task because it will generate a full residue model from the sequence information only, which may deviate at the confident regions in the initial model and consequently may not fit well to the map well. To achieve local refinement with AF2, we modified the input to AF2 in two different ways and compared the results with a regular AF2 run. Thus, we performed three different runs. We used ColabFold (Mirdita *et al*., 2022) on Google Colab to run AF2 because the server allows us to input a customized input.

A regular AF2 run takes an input protein sequence and generates a MSA with MMseqs2 (Mirdita *et al*., 2019, Steinegger & Soding, 2017), which we call here a full MSA, and runs AF2. AF2 simply performs structure prediction and does not consider the initial structure model. This procedure produced five models.

The second run is with the full MSA and a trimmed initial model as a template, where residues with negative DAQ(AA) score, potentially wrongly modelled residue positions, are removed from the initial model. The intention of the trimmed template is to provide a template that only covers confident regions of the initial model. In AF2 trained network models, only two fine-tuned models (model-1 and model-2) use template data (Jumper, Evans, Pritzel, Green, Figurnov, Ronneberger, Tunyasuvunakool, Bates, Zidek, Potapenko, Bridgland, Meyer, Kohl, Ballard, Cowie, Romera-Paredes, Nikolov, Jain, Adler, Back, Petersen, Reiman, Clancy, Zielinski, Steinegger, Pacholska, Berghammer, Bodenstein, *et al*., 2021), therefore this run produces two models by using model-1 and model-2.

The third run is with a trimmed MSA and the trimmed template. In the trimmed MSA, we masked local regions in the full MSA that correspond to the residue with a zero or positive DAQ(AA) score, i.e. confident regions in the initial model. (AF2 with trimmed MSAs + trimmed template). The intention of the trimmed MSA is not to provide MSA information of the confident regions of the model, so that AF2 does not have information to alter the structure of the confident regions. This procedure, too, produces two models. In total, we have nine models for a target protein from the three different AF2 runs. The strategy to trim an MSA and a template was inspired by a work by Heo and Feig, where they attempted to produce multiple conformations of target proteins by controlling an MSA and a template to input (Heo & Feig, 2022).

The nine models were superimposed on the trimmed template, which was then subjected to the Rosetta *relax* protocol (Nivon *et al*., 2013, Conway *et al*., 2014) with the cryo-EM density map. Finally, 18 models, nine models with and without Rosetta relaxation, were evaluated by DAQ(AA) score. We selected the model that has the highest DAQ(AA) score as the final model.

### 2.3. Dataset of PDB entries to refine

We tested the DAQ-refine protocol on thirteen PDB chain models with corresponding EM maps, which were taken from datasets used in the DAQ score paper (Terashi *et al*., 2022). These targets were selected because the initial model of them has mismodelled regions as indicated by negative DAQ(AA) scores.

Six out of the thirteen targets (PDB ID: 6CP3-Y, 6K1H-Z, 7JSN-A, 7JSN-B, 7KSM-C, and 7KSM-D) were PDB entries that have at least two versions of deposited structures, which are available at the wwPDB database (ftp-versioned.wwpdb.org). Moreover, the two versions of the structures are different by more than 1.0 Å Cα RMSD and the first version has a mismodelled region that has a low negative DAQ(AA) score. The second version of the structure has a substantially better DAQ(AA) score in a positive value range, indicating that the error in the model was fixed by the authors. We rebuilt the model from the initial version structure and compared with the second version of the deposited structure. These six targets were named the 2Ver targets. The targets and the file name of PDB files at the PDB’s FTP site are listed in Supplementary Table S1.

The other seven targets (6L54-C, 3J6B-9, 5LC5-N, 6GCS-2, 6GCS-4, 6CV9-A, and 6IQW-E) were selected from protein structure pairs from different PDB entries (and corresponding different EMDB entries) that have over 90% sequence identity to each other; nevertheless have a Cα RMSD of 1.0 Å or higher and have at least four contiguous inconsistent and thus probably misaligned residues between each other. We defined that these misaligned residues are corresponding residues in the two structures that are more than 2.0 Å away from each other and close to different residues when the two models were superimposed. We selected the pairs which one of the pairs had negative DAQ(AA) score for that region. For each structure pair, we refined the structure with a lower DAQ score and examined if the refined structure became close to the higher-scoring counterpart. We call these targets homologous pair (Hom) targets.

There is a possibility that both entries of a Hom target are correct since they are not identical proteins and have been independently deposited to PDB. However, one of the pair has a negative DAQ(AA) score, which usually indicates an incorrect modeling, and our local refinement protocol constantly improved DAQ(AA) when applied to the entry with the lower score. Thus, as we also discussed in the original DAQ paper (Terashi *et al*., 2022), we believe that the entry with the lower score does indeed include wrongly modelled regions. The EMDB and PDB IDs of the entries pair of these targets are provided in Supplementary Table S2. In Supplementary Table S3, we provided the average DAQ(AA) score of each of two structures of the thirteen targets as well as RMSD between them.

## 3. Results and Discussion

### 3.1. Local refinement results with the three AlphaFold protocols

We first show the summary of refined structures built by the three AF2 protocols discussed above, AF2 with a full MSA and no template, AF2 with a full MSA and a trimmed template, and AF2 with a trimmed MSA and a trimmed template. After each model was generated, we ran the Rosetta Relax protocol. For comparison, we also show the results of applying the Rosetta Relax only to the initial models. The results are presented in two metrics, the root-mean-square deviation of Cα atoms (Cα-RMSD) and the high-accuracy version of the global distance test (GDT-HA) (Kopp *et al*., 2007) to the native (i.e. the second version model in the 2Ver targets and the higher scored structure in the Hom targets). GDT-HA is defined as the average fraction of correctly modeled Cα atom positions obtained in each superpositions to the native structure with four distance thresholds, 0.5, 1.0, 2.0, and 4.0 Å. In Table 1, for each protocol, the RMSD and GDT-HA of the best model among generated is shown. Supplementary Table S4 and S5 provide results of all 18 models for each target.

**Table 1.**
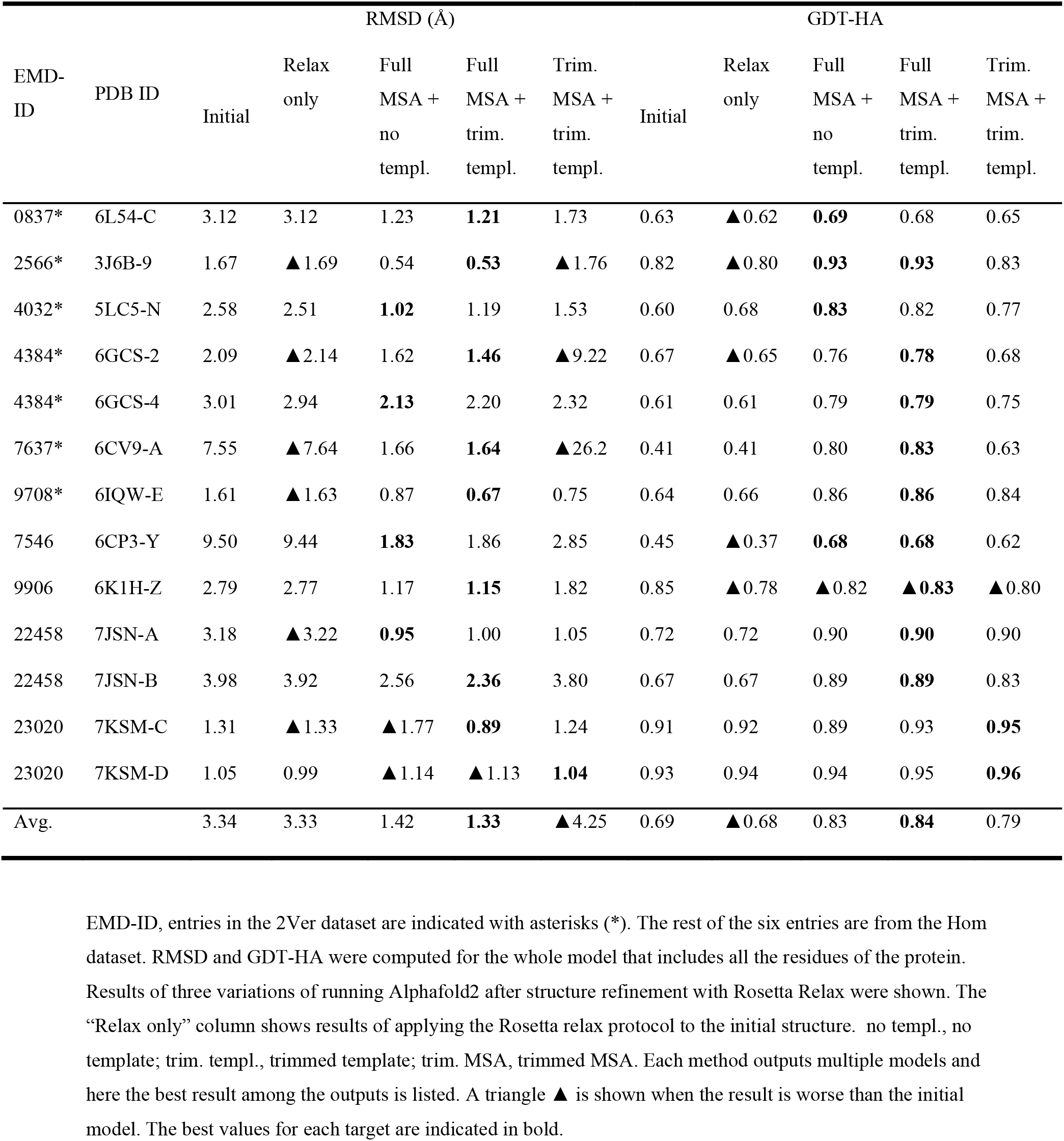
Summary of the structure remodeling with three Alphafold2-based methods.

When Rosetta Relax was applied directly to the initial models, improvement was observed for 6 targets and no change for 1 target (53.8% considering both cases) out of the 13 targets when RMSD was concerned. GDT-HA improved for 4 cases, no change for 4 other cases out of 13. The average GDT-HA went down slightly from 0.69 to 0.68. For both improvements of RMSD and GDT-HA, the Wilcoxon test indicates high p-value, 0.63 and 0.58, respectively. Thus, using only Rosetta Relax did not make statistically significant improvement. This is mainly because misaligned residues in the initial models are difficult to resolve by regular structure refinement.

When we applied three AF2 protocols followed by the Rosetta Relax, the resulting models clearly improved over the simple application of structure relaxation (Table 1). The average RMSD improved from 3.33 Å to 1.42 Å and to 1.33 Å by the AF2 with a full MSA and no template and AF2 with a full MSA and a trimmed template, respectively. The average RMSD went worse for the AF2 with a trimmed MSA and a trimmed template, due to two targets, 6GCS-2 and 6CV9-A, where the resulting RMSD became large values. The RMSD became very large for these two targets because the refined structures had a tail region that was placed in a wrong side of the protein structure. In terms of GDT-HA, all three DAQ-refine protocols improved the initial models except for one target, 6K1H-Z. Comparing the three DAQ-refine protocols, AF2 with a full MSA and a trimmed template performed the best with the smallest average RMSD and the largest average GDT-HA.

To further examine the effect of Rosetta Relax, in Fig. 2, we compared the model accuracy of before and after applying Rosetta Relax. The relaxation improved models for most of the cases. 110 out of 117 models (94.0%) improved and one model stayed the same in terms of RMSD, and 100 models (85.5 %) improved in terms of GDT-HA by the structure relaxation step. For both RMSD and GDT-HA, the Wilcoxon test result indicated the improvements are statistically significant (p-value 1.7e-19 and 2.2e-14). Thus, Rosetta Relax is not helpful when solely used but is effective when combined with the AF2 runs, i.e. the full DAQ-refine protocol.

**Figure 2.**
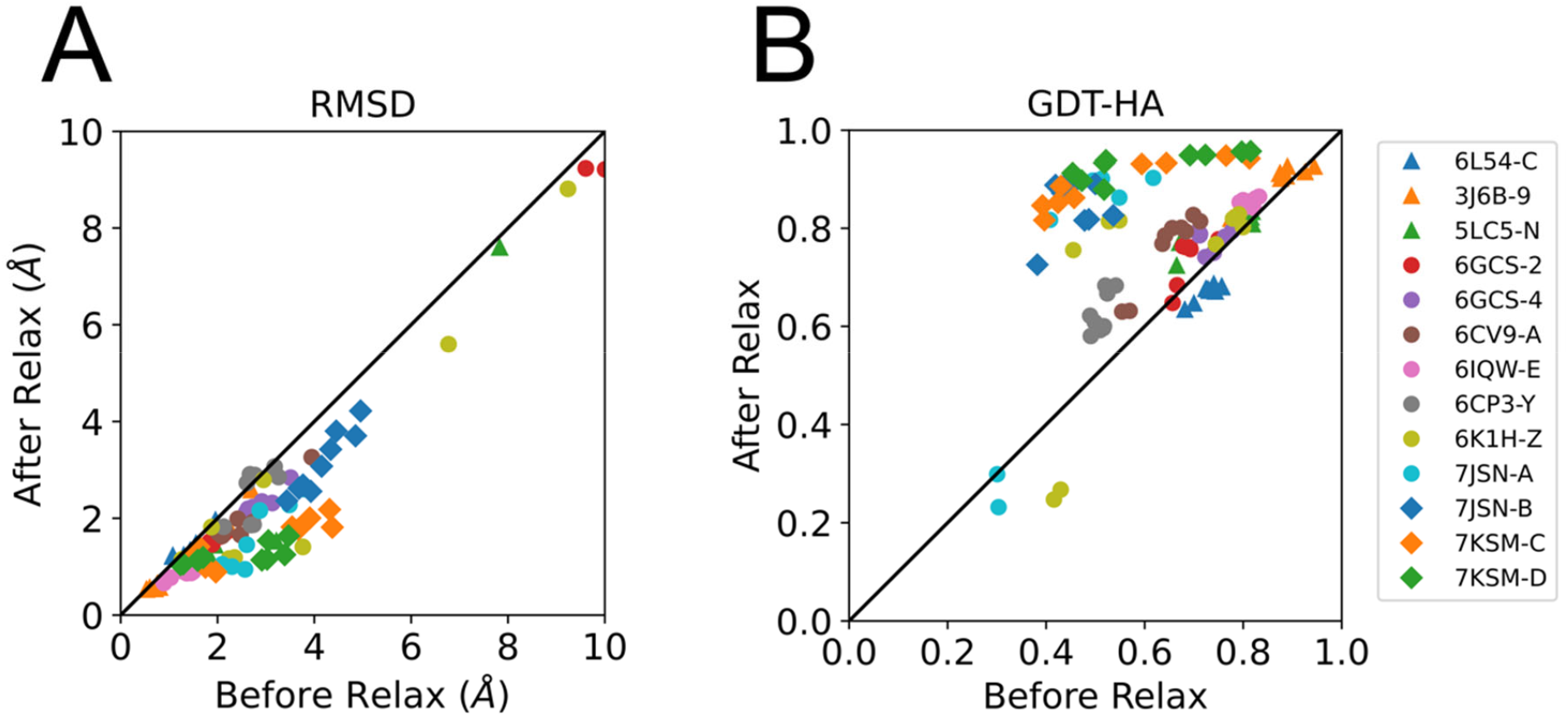
Model quality before and after Rosetta Relax refinement. RMSD and GDT-HA of 9 models for all the 13 targets before and after applying Rosetta Relax structure refinement are shown. **A**, RMSD; **B**, GDT-HA. For RMSD, four models were omitted from the plot that have RMSD values of over 10 Å. They were two models from 6CV9-A, which have before and after RMSD values of (26.0, 26.2) and (35.7, 35.5), respectively; and two models from 7JSN-A, (26.0, 25.6) and (25.4, 25.1).

### 3.2. Model selection with DAQ score

The next question we address is how to select the refined structures out of the 18 models we built. In Fig. 3, we showed GDT-HA for each of the generated 18 models including unrelaxed and relaxed models for each target relative to the DAQ(AA) score. For reference, the initial model and the relaxed initial model from Rosetta Relax are also included in the plots. The same plots instead showing RMSD are provided as Supplementary Fig. S1. The plots show that DAQ(AA) has clear correlation to model quality, GDT-HA. The Pearson’s correlation coefficient between them ranged from 0.10 (6L54-C) to 0.97 (7JSN-B, 7KSM-C, 7KSM-D) with an average of 0.85. Except for 6L54-C, all targets have a correlation coefficient of over 0.85. Therefore, we can use DAQ(AA) as the metric to select one of the most accurate models among those generated.

**Figure 3.**
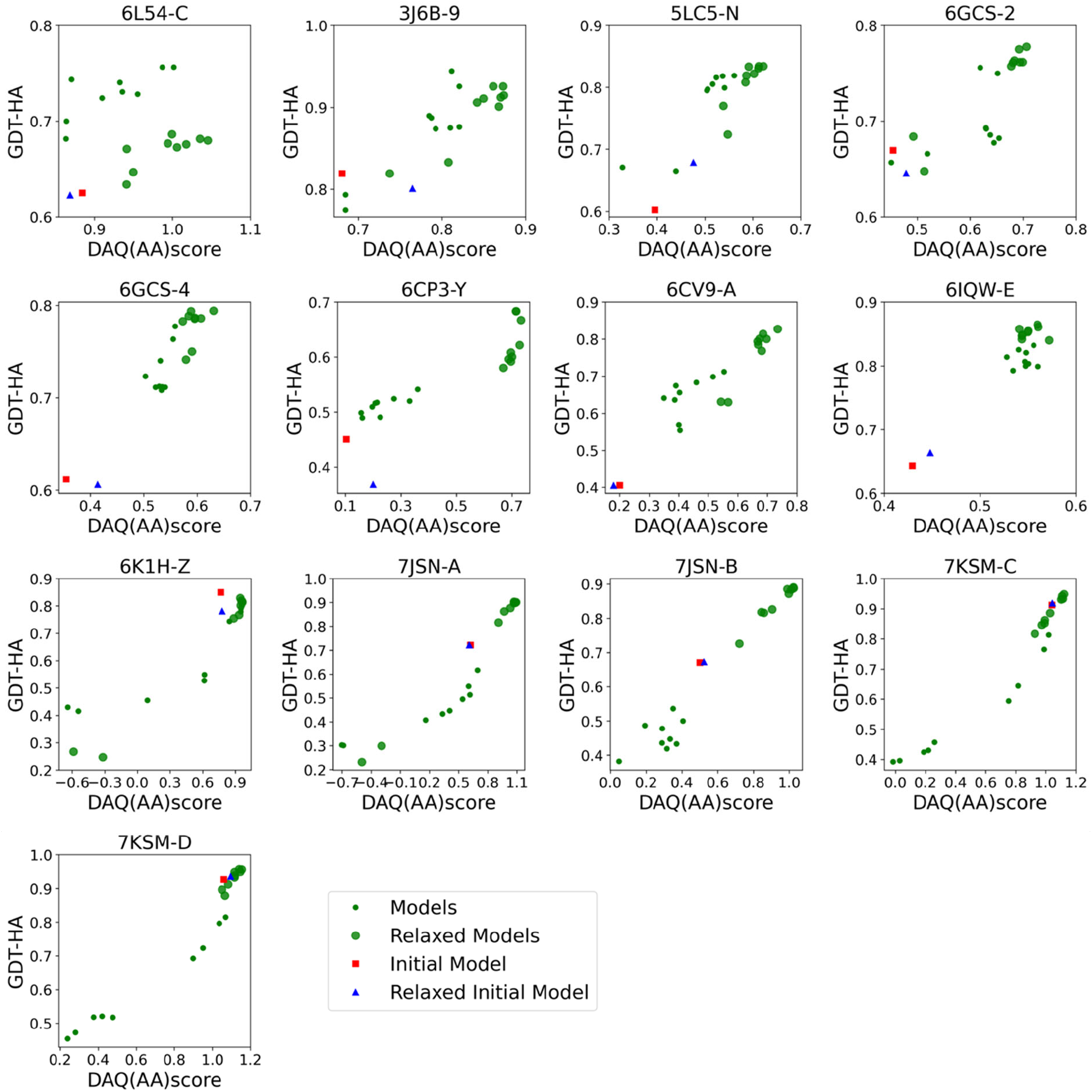
Comparisons of DAQ(AA) score and GDT-HA on 13 targets. DAQ(AA) score of 18 refined structure models with our protocol relative to GDT-HA. Small green circles represent the refined models without the Rosetta relaxation protocol. Small green circles represent the refined models without the Rosetta relaxation protocol. Large green circles represent the refined models after the Rosetta relaxation protocol. Initial models and relaxed initial models by Rosetta relaxation are shown by red square and blue triangle, respectively.

Fig. 4 shows actual model selection results using DAQ(AA). The model with the largest DAQ(AA) improved over the initial model in terms of both RMSD and GDT-HA (Fig. 4A, 4B). There was one target for which the largest DAQ(AA) model had a tie with the initial model in RMSD and was worse in terms of GDT-HA than the initial model. The average RMSD improvement is 1.96 Å, while the improved margin for GDT-HA was on average 0.15 from the initial models. For both improvements of RMSD and GDT-HA, the Wilcoxon test indicates statistical significance with low p-values, 4.9e-4 and 1.2e-3.

**Figure 4.**
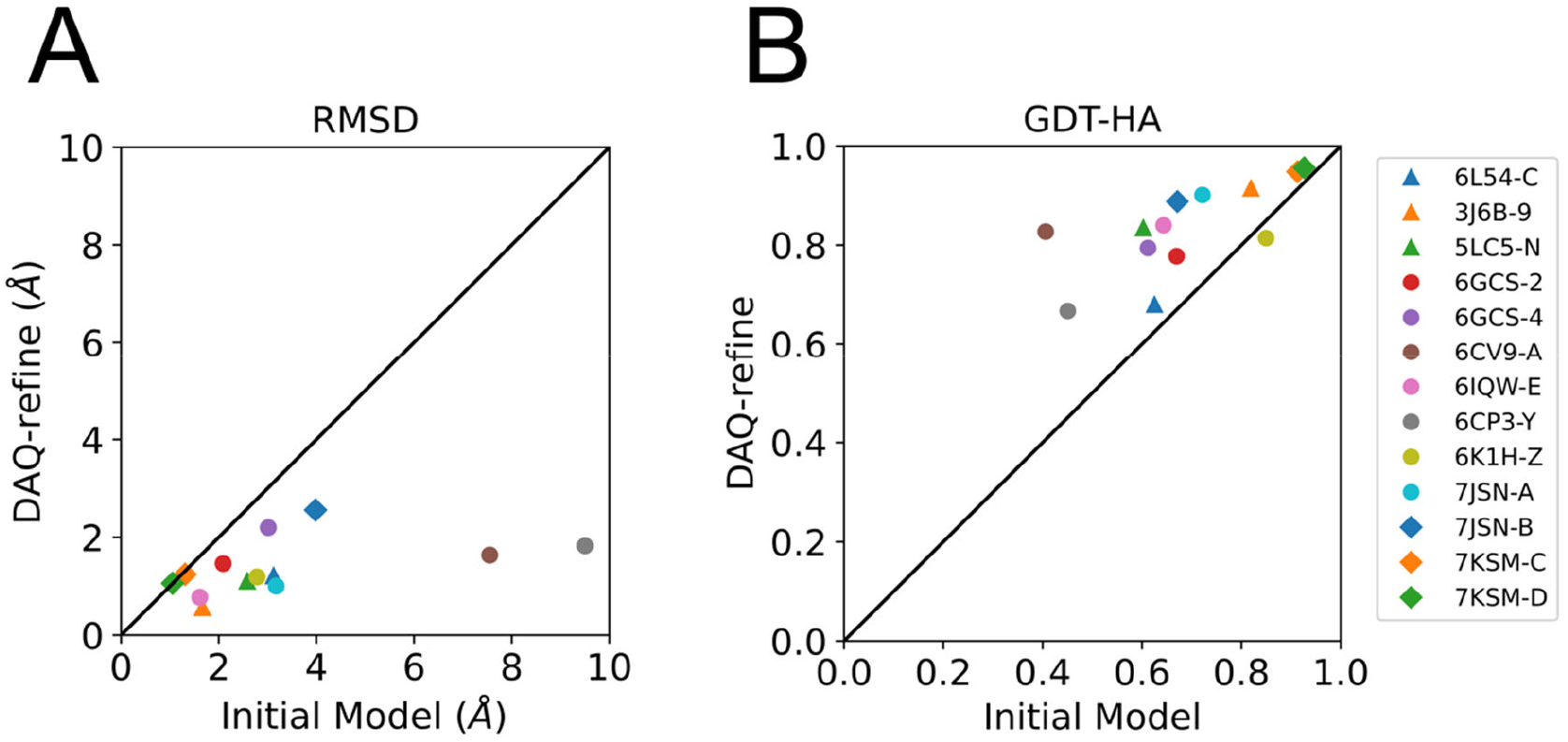
Model selection with DAQ(AA). Out of 18 models generated for each target, the one with the highest DAQ(AA) was selected. The selected models were compared with the initial model in terms of **A**, RMSD; **B**, GDT-HA.

### 3.3. Comparison with other structure refinement methods

We further compared the DAQ-refine protocol with four other existing refinement methods, namely: Molecular Dynamics Flexible Fitting (MDFF) with a g-scale of 0.5 (Singharoy *et al*., 2016, McGreevy *et al*., 2016), the Rosetta relax protocol (Nivon *et al*., 2013, Conway *et al*., 2014), *phenix*.*real_space_refine* (Afonine *et al*., 2013), and *phenix*.*dock_and_build* (Steinegger & Soding, 2017). MDFF performs structure refinement with molecular dynamics under the constraint of an input density map. *phenix*.*real_space_refine* performs gradient-driven minimization of the target function that combines the fit of the model to the map and the restraints of the protein structure, such as bond lengths and angles. *phenix*.*dock_and_build* first runs AF2 to predict the tertiary structure of the target protein and then splits it into reliable domains based on the pLDDT score, then performs docking and rebuilding iteratively for each domain in the map.

Fig. 5 compares RMSD and GDT-HA of refined structures for the 13 targets by our DAQ-refine protocol and the results by the four existing methods. Numerical values of the data are provided in Supplementary Table S6. It should be noted that *dock_and_build* constructs models for only the confident regions as shown in Supplementary Table S7. On average, models by *dock_and_build* cover 89% of amino acids in the target proteins and the rests were not modelled. Therefore, models generated by *dock_and_build* tend to have low (good) RMSDs but low coverage that results in low GDT-HA.

**Figure 5.**
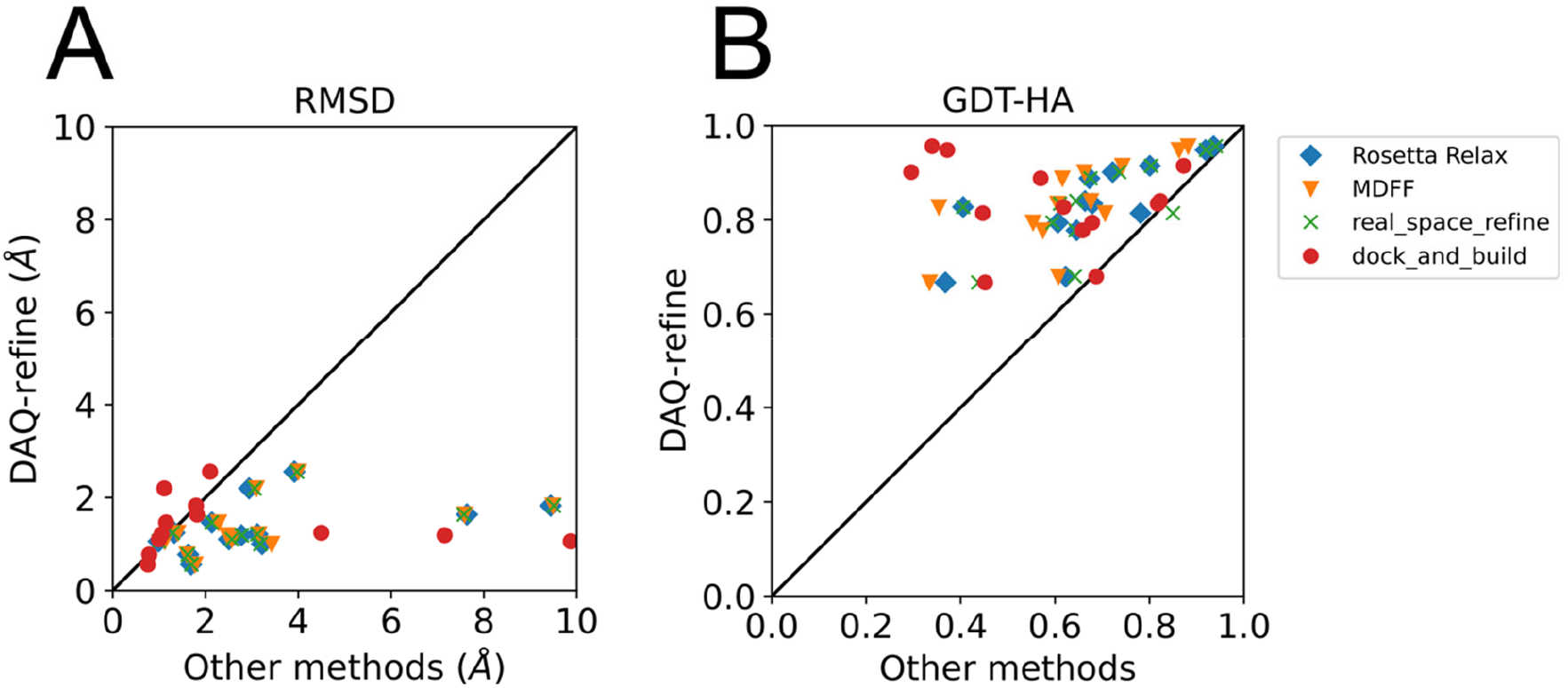
Comparison of DAQ-refine with four other existing methods. For each of the 13 targets the model with the highest score was selected among models generated by each method. Blue diamonds, Rosetta Relax; orange triangles, MDFF; green crosses, *phenix*.*real_space_refine*; and red circles, *phenix*.*dock_and_build*. The *phenix*.*dock_and_build* protocol starts by predicting the structure of the target protein by AF2. For the other refinement methods started from the initial protein model. For our protocol, the model with the highest DAQ(AA) score was selected. **A**, comparison is made in terms of RMSD to the native structure. There was one model generated by *phenix*.*dock_and_build* that has a large RMSD of 25.6 Å and was not included in this plot. **B**, GDT-HA.

Our protocol achieved a lower RMSD for all the targets when compared with Rosetta Relax, MDFF, and *phenix*.*real_space_refine* (Fig. 5A). When compared with models from *phenix*.*dock_and_build*, DAQ-refine had a larger RMSD than *phenix*.*dock_and_build* for 6 targets (6L54-C, 5LC5-N, 6GCS-2, 6GCS-4 6CP3-Y, and 7JSN-B) models, but this is mainly because *phenix*.*dock_and_build* did not model all of the residues in the proteins, i.e. their models are shorter than the native structure. When GDT-HA is considered, DAQ-refine showed a higher value for all the targets than Rosetta Relax and MDFF. Against *phenix*.*real_space_refine* and *dock_and_build*, our protocol showed a higher GDT-HA for 12 (92.3%) targets.

### 3.4. Case Studies

The following sections discuss four case studies that illustrate how DAQ-refine has improved initial models.

#### 3.4.1. Case Study 1, PDB entry 7JSN-A (EMD-22458)

In this example, we refined the first version model of cGMP-specific 3’,5’-cyclic GMP phosphodiesterase (cGMP phosphodiesterase 6 subunit PDB ID: 7JSN-B (Gao *et al*., 2020)) (Fig. 6A). The model was built from a 3.2 Å EM map (EMD-22458). PDB 7JSN has two versions of structures in PDB. The first model was released from PDB on October 21, 2020, which was later revised on March 31, 2021 (Gao *et al*., 2021). In the initial model, which is shown on the left panel of Fig. 6A, there are several regions colored in magenta where large deviations over 3.0 Å were observed when compared with the revised version model. DAQ(AA) score clearly indicates low values for these inconsistent regions as shown in red in the second panel from the left.

**Figure 6.**
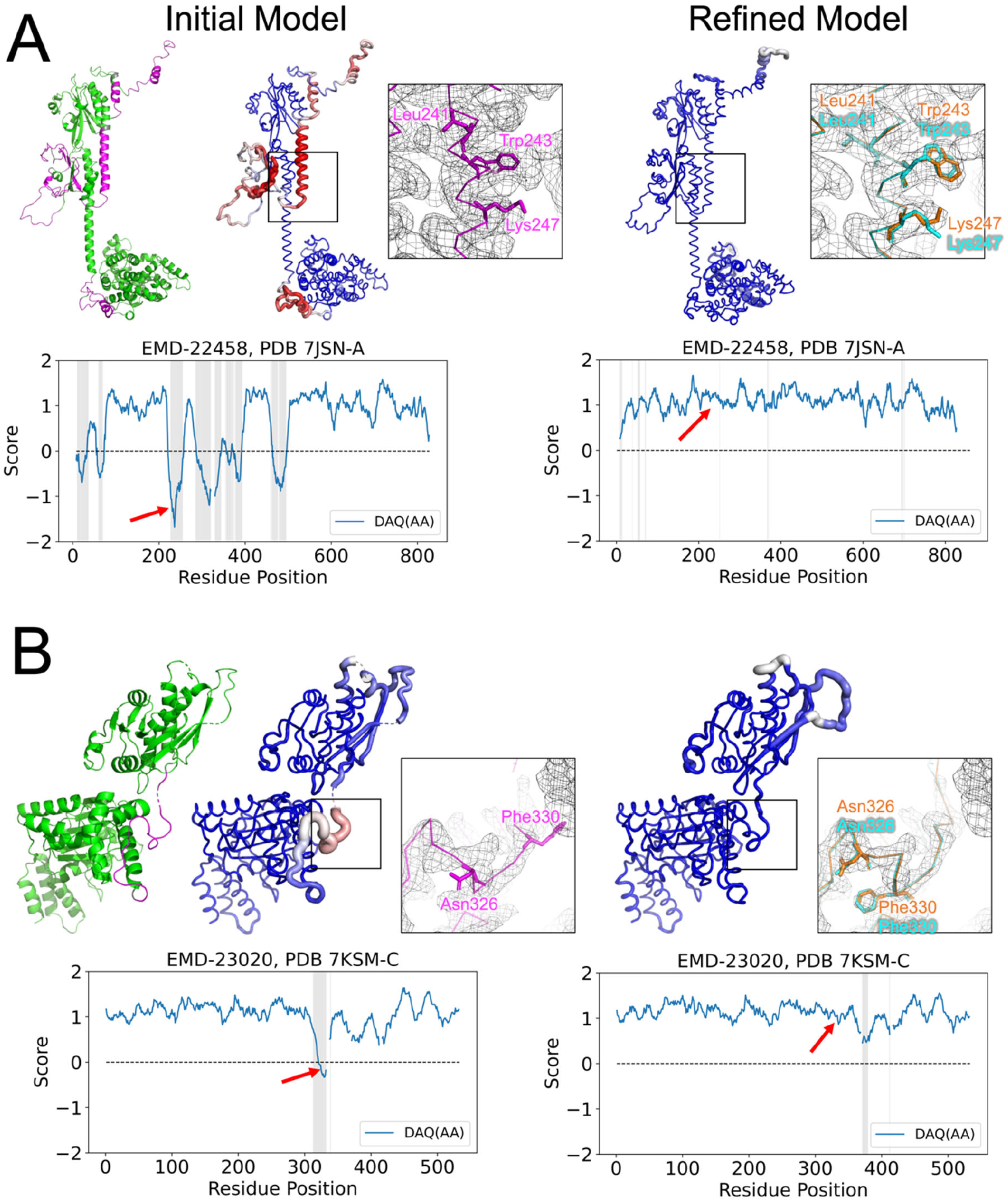
Analysis of initial and refined models of two 2Ver targets by DAQ(AA) score. Left: the initial model (the first version of the PDB entry). Colors in the chain reflects the deviation of Ca atom positions from the native structure (the revised version of the entry). Colors are scaled from green (deviation < 1.0 Å) to magenta (deviation > 3.0 Å). Middle and Right: the initial and the refined models colored by DAQ(AA) score. The color is scaled from red (DAQ(AA) < −1.0) to blue (DAQ(AA) > 1.0). The radius of the chain tube is thicker if the region has a low DAQ(AA) score. In magnified boxes shown are model regions with a low DAQ(AA) score. The initial, the refined, and the revised version model are indicated in magenta, cyan, and orange models, respectively. Surface meshes represent the EM map at the author’s recommended contour level. Plots show DAQ(AA) scores along the sequence position. Gray in the plot represents residue positions where the deviation of the Ca atom position between the model and the native (the revised version structure) is larger than 3.0 Å. A. PDB entry 7JSN chain A, which was built from the EM map EMD-22458. A region with Leu241, Trp243, and Ly247, which has a residue shift in sequence assignment in the initial model is magnified in squares. The corresponding position in the plot is indicated by a red arrow. In the refined model, orange are residue conformations in the native (the second version model) and the cyan residues are refined residues by DAQ-refine. B. PDB entry 7KSM chain C with the EM map (EMD-23020) from which the chain structure was built. A region with a residue shift in the initial model, which include Asn326 and Phe330, is magnified in squares.

**Figure 6.**
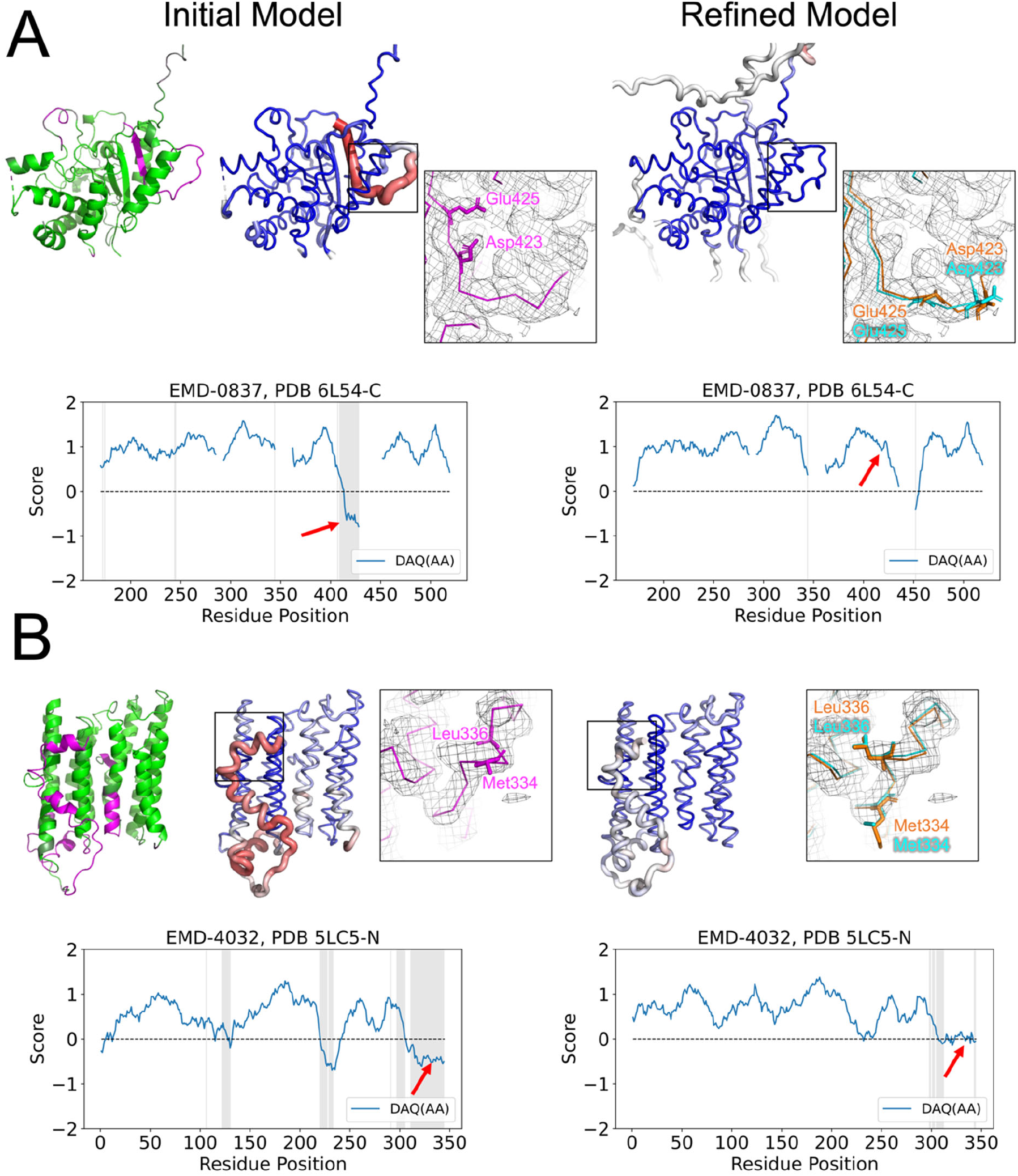
Analysis of initial and refined models of two Hom targets by DAQ(AA) score. **A**. The PDB entry 6L54-C built from the EM map EMD-0837 had regions that are inconsistent with the other protein model, PDB: 6Z3R-C from the EM map EMD-11063. The region is shown in magenta in the leftmost structure and detected as negative DAQ(AA) as shown red with a thick tube in the structure model placed the second from the left. The refined model is shown on the right. It has overall positive score (blue). The two squares provide magnified view of residues in the inconsistent region before and after the refinement by DAQ-refine. The DAQ score plots have gaps at missing residues in the initial model (6L54-C). In magnifying square windows, Asp423 and Glu425 in pink are conformations of these two residues in the initial model whereas those in cyan are results of DAQ-refine. Those in orange are conformations in 6Z3R-C. **B**. Local refinement of the PDB entry 5LC5 chain N, which was built from the EM map EMD-4032 in comparison with PDB: 6ZKM-N, which was derived from EMD-11254. Magnifying squares highlight Met334 and Leu336 in an inconsistent region between the two entries.

The refined structure is shown on the right in Fig. 6A. DAQ(AA) score of the refined model is much improved, positive values for all the residues, as indicated in blue in the model. The change of the scores along the residue positions is clear in the two plots shown below the model structure images. In the initial model, inconsistent regions to the native (shown in gray) had low, negative DAQ(AA) score while the refined model have all positive DAQ(AA) values along the chain. One of the remodeled regions is highlighted in squares. In the initial model, residues including Leu241, Trp243, and Lys247 do not fit well to the density. The problem of this region is that the assigned sequence was shifted along the helix. On the other hand, in the refined model shown on the right, the revised conformation of the three residues (cyan) agree well to the native structure except for the direction of the tip of Lys247. After the local refinement, overall model accuracy was improved from 3.18 to 1.00 Å in Cα-RMSD and from 0.72 to 0.90 in GDT-HA.

#### 3.4.2. Case Study 2, PDB entry 7KSM-C (EMD-23020)

The second example is a model of human mitochondrial AAA+ protein LONP1. The first version model (PDB ID: 7KSM (Shin *et al*., 2021)) was released from PDB on December 2, 2020 and then it was revised on June 15, 2022. As shown in Fig. 6B left plots, there is an inconsistent region (Ile312-Glu339, gray shade in the score plot and shown as magenta in the left-most model) between the first and revised version model, where DAQ(AA) exhibited negative values as indicated with a thick tube in red. As highlighted in the square, side-chains of Phe330 and Asn326 in the first version model (magenta) were not covered well by the map density. The main problem of this region is that the sequence assignment was shifted along the main-chain conformation, which resulted in the unnatural side-chain conformation in the density. In the refined model (the right panel) now DAQ(AA) for these two amino acid residues improved to positive values and they are fitted well in the map. After the refinement, overall model accuracy was improved from 1.31 in 1.24 Å of Cα-RMSD and 0.91 to 0.95 in GDT-HA.

#### 3.4.3. Case Study 3, PDB entry 6L54-C (EMD-0837)

The next two examples are taken from the Hom targets. Fig. 7A is a pair of models from the protein SMG9. One is PDB: 6L54 chain C, which was modelled from the EM map EMD-0837, determined at a 3.43 Å resolution (Zhu *et al*., 2019) and the other is PDB: 6Z3R chain C, built from the EM map EMD-11063, 2.97 Å resolution (Langer *et al*., 2020). These two protein models have 100% identical sequences, yet their structures deviate by an RMSD of 3.1 Å. In 6L54 chain C, Thr405 through Leu428 has a negative DAQ(AA) score (shown red in the second model from the left), indicating likely a shift in the sequence assignment. By close inspection, it is observed that this region has implausible side-chain packing. For example, side chains of Asp423 and Glu425 are directed into the hydrophobic core of the protein in 6L54 (magenta, highlighted in the magnifying square). But in 6Z3R, these residues are exposed to solvent (the orange model, highlighted in square on the right), which would be more appropriate. 6L54 chain C has a total DAQ(AA) score of 272.4, lower than 6Z3R chain C’s DAQ(AA) score of 308.2. Therefore, we used 6L54 chain C as the initial model and refined it to see if the structure gets closer to the 6Z3R chain C as the native structure. As shown in the right column in Fig. 7A, DAQ-refine corrected the inconsistent region and improved the DAQ(AA) score. After local refinement, Asp243 and Glu425 in the 6L54 chain C (cyan in the magnifying square on the right) took similar conformations to those in 6Z3R chain C (orange). Reflecting this change, DAQ(AA) score improved to positive values as shown in the plots.

#### 3.4.4. Case Study 4, PDB entry 5LC5-N (EMD-4032)

The last example (Fig. 7B) is a pair of structures of NADH-ubiquinone oxidoreductase chain 2, the PDB entry 5LC5 chain N (EMD-4032, 4.35Å resolution (Zhu *et al*., 2016)) and PDB: 6ZKM chain N (EMD-11254, 2.8 Å resolution (Kampjut & Sazanov, 2020)). These two protein models have a high sequence identity of 91.9% but have a structure deviation of a 2.6 Å RMSD. There are three inconsistent regions between the two PDB models, which are indicated in magenta in the leftmost structure model and in gray shades in the DAQ(AA) plot on the left. These regions are identified as having a low, negative DAQ(AA) score. Applying the refinement protocol to 5LC5 chain N modified the sequence assignment of this region, and improved DAQ(AA) from negative values to positive as shown in the two plots (the positions indicated by red arrows). The refined model of 5LC5-N has now the identical sequence assignment as 6ZKM-N. To illustrate the refined model, we show two residues, Leu336 and Met334. In the initial model, 5LC5-N, these two hydrophobic residues are facing solvent and do not fit well to the density. The refined structure (cyan in the magnified square on the right) now has the two residues in the same conformations as 6ZKM-N, where they face the interior of the protein having hydrophobic interactions, which would be more reasonable.

In all these four examples, DAQ(AA) score detects inconsistent regions between two models compared with a negative score in one of the models, and those regions were improved to a positive DAQ(AA) score by the DAQ-refine protocol.

## 4. Summary

As more protein structure models built from high-resolution EM maps become available, accurate model evaluation and suitable local refinement methods become more important. In this work, we presented a protein structure local refinement protocol, DAQ-refine, which uses the local model quality evaluation with DAQ score to detect potential local incorrect regions and rebuilds them with AlphaFold2. To reflect the local quality data by DAQ score in the refinement step by AlphaFold2, we introduced a trimmed template model and trimmed MSAs. Trimmed input data allows AlphaFold2 to focus on building incorrect regions while keeping correct regions almost intact. Our protocol generates a series of different models using different AlphaFold2 settings. We demonstrated that DAQ(AA) has a substantial correlation to the quality of the models and is able to select good models among generated.

## Availability

The DAQ program is freely available for academic use via GitHub, https://github.com/kiharalab/DAQ. In addition, the DAQ program is available to run on a Google Colab notebook at https://bit.ly/daq-score and https://github.com/kiharalab/DAQ/blob/main/DAQ_Score.ipynb. DAQ-refine including the modified ColabFold that can use both trimmed MSAs and a trimmed template is available at https://bit.ly/DAQ-Refine and https://github.com/kiharalab/DAQ-Refine. This colab notebook provides the instructions of the DAQ-refine and tools for generating trimmed template and trimmed MSAs based on the results of DAQ program.

## Acknowledgements

We thank Charles Christoffer for testing the DAQ-refine program on Google Colab and also for proofreading the manuscript. This work was partly supported by the National Institutes of Health (R01GM133840, R01GM123055, and 3R01GM133840-02S1) and the National Science Foundation (CMMI1825941, MCB1925643, DBI2146026, and DBI2003635).

