## Supplementary File 1 for "Protein Model Refinement for Cryo-EM Maps Using DAQ score"

**Supplementary Table S1. Six EMDB entries and protein chains in the 2Ver targets.**

| EMID | resolution (Å) | PDB ID | CIF file of the first version | CIF file of the revised version |
|---|---|---|---|---|
| 7546 | 3.8 | 6CP3-Y | pdb_00006cp3_xyz_v1.cif | pdb_00006cp3_xyz_v2.cif |
| 9906 | 3.52 | 6K1H-Z | pdb_00006k1h_xyz_v1.cif | pdb_00006k1h_xyz_v2.cif |
| 22458 | 3.2 | 7JSN-A | pdb_00007jsn_xyz_v1.cif | pdb_00007jsn_xyz_v2.cif |
| 22458 | 3.2 | 7JSN-B | pdb_00007jsn_xyz_v1.cif | pdb_00007jsn_xyz_v2.cif |
| 23020 | 3.2 | 7KSM-C | pdb_00007ksm_xyz_v1.cif | pdb_00007ksm_xyz_v2.cif |
| 23020 | 3.2 | 7KSM-D | pdb_00007ksm_xyz_v1.cif | pdb_00007ksm_xyz_v2.cif |

Each target has two versions of the protein structure. The fifth and sixth columns show the file names of the deposited models in the wwPDB database (ftp-versioned.wwpdb.org).

**Supplementary Table S2. Seven protein chain pairs in Homologous pair targets.**

| | Initial Model | | | Native Model | |
|---|---|---|---|---|---|
| EMID | Resolution (Å) | PDB ID | EMID | Resolution (Å) | PDB-ID |
| 0837 | 3.43 | 6L54-C | 11063 | 2.97 | 6Z3R-C |
| 2566 | 3.2 | 3J6B-9 | 3556 | 3.2 | 5MRF-9 |
| 4032 | 4.35 | 5LC5-N | 11254 | 2.8 | 6ZKM-N |
| 4384 | 4.32 | 6GCS-2 | 4872 | 3.3 | 6RFQ-2 |
| 4384 | 4.32 | 6GCS-4 | 4872 | 3.3 | 6RFQ-4 |
| 7637 | 3.8 | 6CV9-A | 11674 | 3.62 | 7A6U-A |
| 9708 | 3.35 | 6IQW-E | 9255 | 3.1 | 6MUT-E |

Each target has two protein chains. For each target, the initial model has a lower DAQ score than the native model and used as the starting model for remodeling. The native model is the structure, which the initial model and remodeled structure were compared against. The average DAQ(AA) score and RMSD between the initial model and native models are shown in Supplementary Table S3.

**Supplementary Table S3. Model quality and DAQ(AA) score of the initial model and the native models of the targets. (in a separate Excel file).**

Each target has two structures. The initial model is the model to which the remodeling protocols were applied. The native model is the other model, which was considered as the correct structure of the protein of the initial model.

**Supplementary Table S4. RMSD of all remodelled structures by three AF2-based protocols (in a separate Excel file).**

For each of the 13 targets, RMSD of the models built by the three AF2-based protocols are shown. RMSD of before and after Rosetta Relax refinement are shown.

**Supplementary Table S5. GDT-HA of all remodelled structures by three AF2-based protocols (in a separate Excel file).**

For each of the 13 targets, GDT-HA of the models built by the three AF2-based protocols are shown. GDT-HA of before and after Rosetta Relax refinement are shown.

**Supplementary Table S6. RMSD and GDT-HA of models built by four existing remodelling protocols (in a separate Excel file).**

Numerical values of remodelled structures of the 13 targets by the four methods, Rosetta Relax, MDFF, phenix.real_space_refine, and phenix.dock_and_build are shown. These data are used to plot Fig. 5.

**Supplementary Table S7. Coverage of models generated by phenix.dock_and_build (in a separate Excel file).**

A model built by phenix.dock_and_build does not necessarily include all the residues in the target because the procedure starts from fitting reliable regions of a AF2 model into the density map. In this table, the fraction of modelled residues in each target is shown.

**Supplementary Figure 1. Comparisons of DAQ(AA) score and Ca-RMSD of models for the 13 targets.**

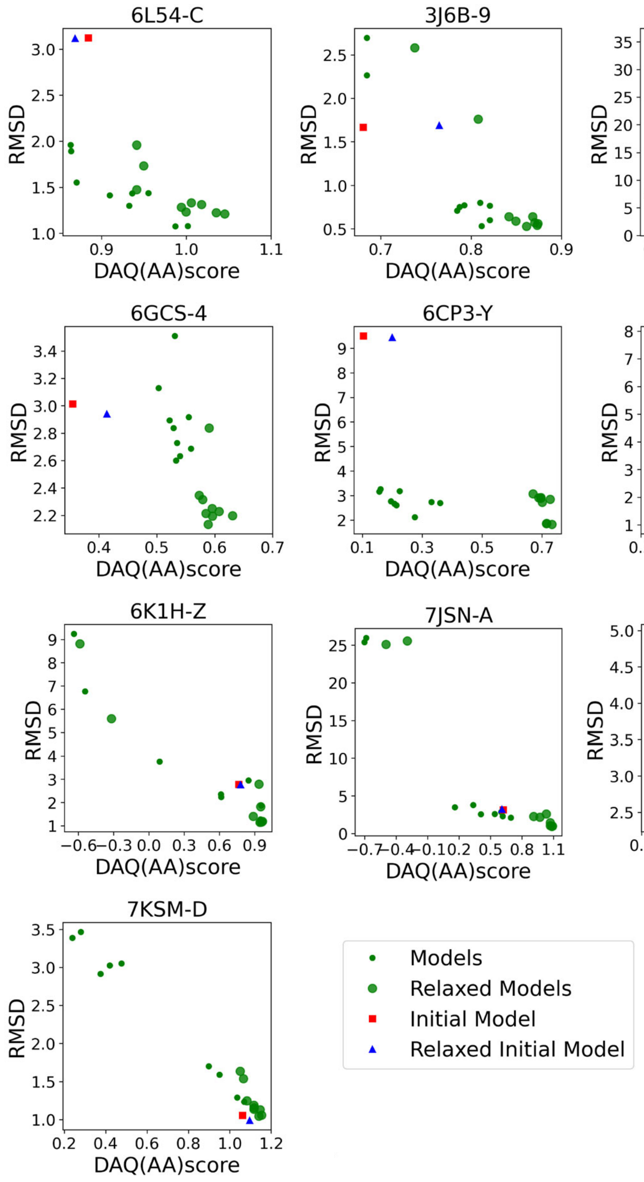


Small green circles represent the refined models without the Rosetta relaxation protocol. Large green circles represent the refined models after the Rosetta relaxation protocol. Initial models and relaxed initial models by Rosetta relaxation are shown by red square and blue triangle, respectively.
